# Diffuse predictions stabilize and reshape the neural code during working memory encoding

**DOI:** 10.64898/2026.02.23.707359

**Authors:** Nursena Ataseven, Șahcan Özdemir, Wouter Kruijne, Daniel Schneider, Elkan G. Akyürek

**Author notes:** Equal senior authorship (Wouter Kruijne, Daniel Schneider, Elkan G. Akyürek).

## Abstract

Predictions can alter working memory (WM) representations. However, its effects may have been mischaracterized due to the use of *precise* predictions in previous experiments, where exact properties of upcoming memory items are cued in advance. Here we investigated a more ecologically valid scenario, in which we assessed the impact of *diffuse* predictions, where advance cues provided only partial knowledge about the targets. To investigate the resultant nature of the target representations in WM, we performed a series of multivariate analyses of EEG data. Forty participants judged whether a probe grating was rotated clockwise or counterclockwise relative to a memorized orientation, which was either predictable or unpredictable. Each memory item was preceded by a central color cue (red, green, or blue). In half of the trials, two of these (predictive) colors cued two non-overlapping 90° segments of orientations that the grating was sampled from. Thus, participants knew the range of possible orientations of these items, but not their exact orientation. In the other half of the trials, a third (non-predictive) color was presented, signaling that the item could have any possible orientation. Behavioral results revealed higher accuracy for predictable items, with systematic biases toward the center of the cued segment. EEG results revealed equally successful decoding of orientation for both predictable and unpredictable items during memory encoding. However, cross-condition decoding was significantly weaker than within-condition decoding, suggesting that the encoding format changed between conditions. Representational similarity analysis showed higher similarity between predictable items, with a representational bias towards the cued segment. Covariance matrices showed lower variance for predictable items while the representational space of predictable items was shrunk. These effects were absent during the maintenance phase. Together, our findings suggest that diffuse predictions alter the geometric layout of the neural representations and stabilize the neural code during WM encoding.

## Introduction

The human brain could be viewed as a prediction machine (Clark, 2013). It constantly generates internal models to predict incoming sensory information, and updates these internal models based on experience to eventually minimize prediction errors (Clark, 2013, 2015; Friston, 2005, 2010). Therefore, perception and cognition are not merely shaped by incoming sensory information, but reflect an integration of incoming information with internal predictions (Friston, 2005, 2010; Körding & Wolpert, 2004; Lee & Mumford, 2003; Rao & Ballard, 1999). A growing body of neuroimaging research has provided evidence for this, by showing that predictions not only benefit behavioral performance but also influence neural activity and representations (Agres et al., 2018; Beck et al., 2014; Blank & Davis, 2016; Bollinger et al., 2010; Brady et al., 2009; Eissa & Kilpatrick, 2023; Giesbrecht et al., 2006; González-García & He, 2021; Katzner et al., 2012; Kok et al., 2012; Kumar et al., 2017; Padgaonkar et al., 2017; Ravizza, 2025; Summerfield & Mangels, 2006; Yan et al., 2023; Yon et al., 2018).

For example, Kok et al. (2012) investigated the effect of prediction on neural representations by using a working memory (WM) task. Participants memorized an orientation grating and compared its orientation or frequency to a probe grating. An auditory cue that preceded the memory grating predicted its orientation with 75% validity. They showed that when the grating’s orientation matched the cued prediction, its decodability in the visual cortex, as measured with fMRI, was enhanced. The authors concluded that predictability of the upcoming information sharpened the neural representations.

The literature on prediction has presented different types of predicting the identity of upcoming information. Examples may be precise predictions, where a cue directly predicts a stimulus (Kok et al., 2012; Yan et al., 2023; Yon et al., 2018) or probabilistic predictions, where information is predicted based on a learned probabilistic prior distribution of outcomes (Barber et al., 2003; Brady et al., 2009; Eissa & Kilpatrick, 2023). Here, we propose another type of deterministic prediction with high ecological validity; *diffuse predictions*. These predictions inform about the upcoming information in an inexact manner, spanning a broad range of possibilities with equal likelihood. Diffuse predictions provide uniform priors that could be built from broad experiences in the world, as in a Bayesian framework (Friston, 2005, 2010; Körding & Wolpert, 2004; Lee & Mumford, 2003; Rao & Ballard, 1999). For instance, when you are in the Italian food section in a grocery store, you might ‘diffusely’ expect to see products like pasta and tomato sauce over corn and jalapeño even though you may not have specific expectations of the products. Such diffuse predictions may require a representational shift that is different from precise predictions to allow for a variety of predictions without a specific likelihood. In this experiment, we manipulated the availability of diffuse predictions and investigated whether and how they affect neural representations during information encoding and working memory (WM) maintenance.

In this paper, we conducted an in depth investigation of the neural representations by analyzing neural variance and structural geometry of the same stimuli either when they are or aren’t predicted. To do this, we explicitly taught participants cues that either predicted the upcoming memory item (an orientation grating) or did not provide any information. Predictions were diffuse, in that they were associated with a range of orientations from which the memory item could be selected with equal likelihood. We recorded their EEG, and investigated how diffuse predictions affected neural item representations during encoding as well as during maintenance. To better observe representations during WM maintenance, we showed an impulse stimulus in the retention interval and analyzed the dynamic EEG impulse response (Wolff et al., 2015, 2017). We hypothesized that the predictions will alter the shape of the representations such that the representational change will reflect an influence of the predicted orientation range. As sharpening of the neural representations might be specific to precise predictions, we do not expect such an effect of the diffuse predictions in the current study.

## Results

EEG data were recorded from forty participants who performed a delayed comparison task (Figure 1A), in which they memorized an orientation grating and later judged whether a probe grating was rotated clockwise (CW) or counterclockwise (CCW) in comparison. Before each memory item, a color cue (red, green, or blue) was shown. On half the trials, two predictive cue colors indicated distinct 90° orientation segments (referred to as templates) from which the memory item was going to be drawn. On the other half, a third nonpredictive color was shown, which signaled that the memory item could have any orientation. Participants learned these associations before the experiment.

**Figure 1.**
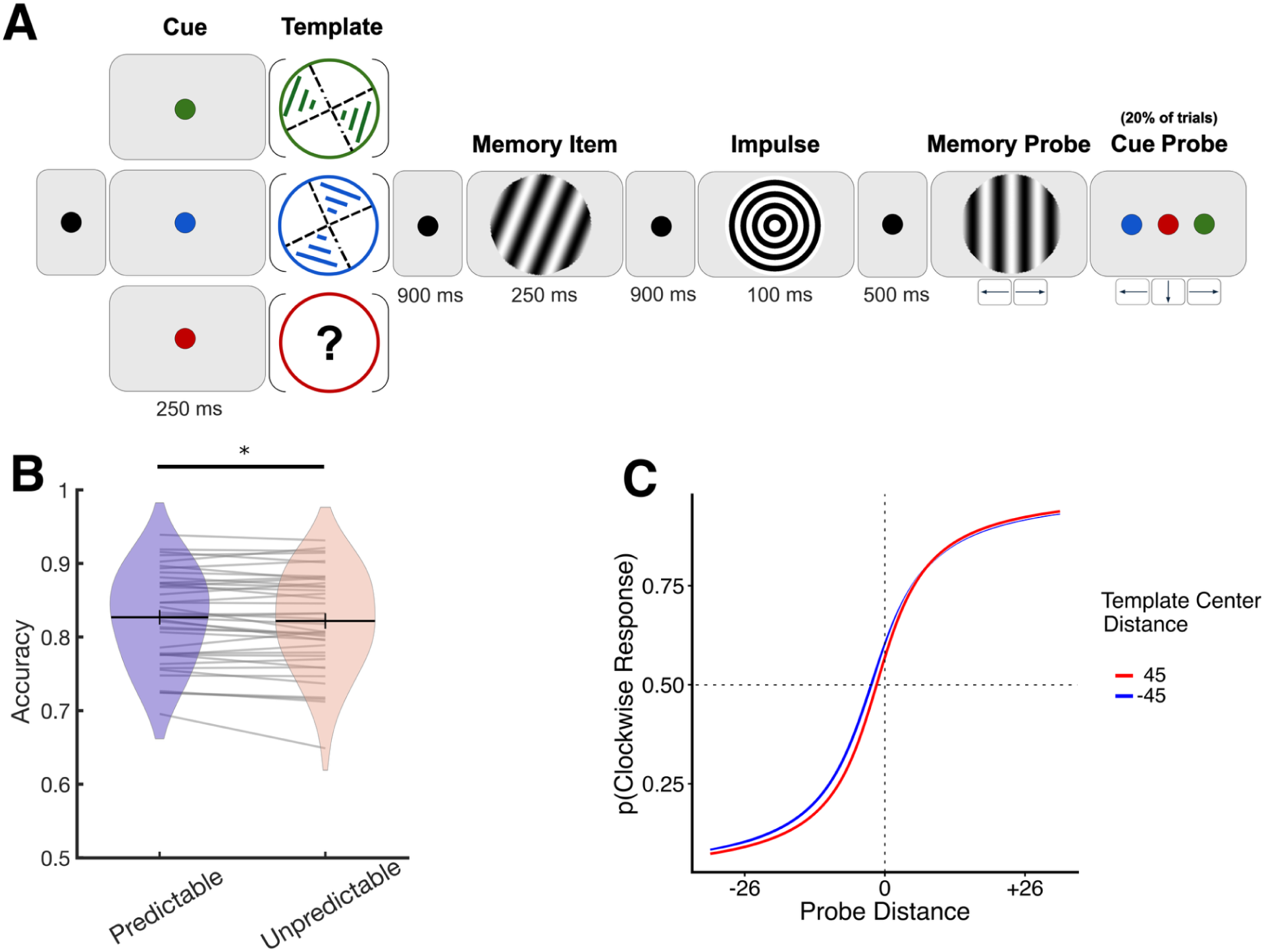
Experimental procedure and behavioral results. **(A)** Depiction of a trial. Participants first saw a cue that was either associated with a prediction (visualized as green and blue) or not (red). Later, they memorized an item that conformed to the cued prediction. They judged whether the memory probe was CW or CCW relative to the memorized item. The impulse was shown in the retention interval to assess the representation in WM during maintenance. The cue color itself was probed in 20% of the trials to keep participants attending to the cue display. **(B)** Violin plot that shows the mean accuracy per condition averaged over probe distances. Gray lines depict individual participant scores. (* *p <* 0.05) **(C)** Probability of giving a CW response as a function of probe distance and template center distance. The lines show the binomial regression fit evaluated at -45 and +45 degrees of template.

### Predictive cues improve memory performance

We investigated the effect of memory item predictability on the probability of correct responses using generalized linear mixed-effects models (GLMM; see Methods). A likelihood ratio test revealed that a model including *memory item predictability* was preferred over an intercept-only model, χ²(1) = 4.20, p = 0.040. The model fit showed a subtle behavioral benefit for trials with predictive cues, β = 0.06, CI = [0.003, 0.116] (Figure 1B).

### Responses were biased towards the center of prediction

To measure how memory item predictability benefited performance exactly, we next explored whether participants’ responses were biased with respect to the template center. To this end, we fit a GLMM to data from the predictable condition, including terms for probe distance (current orientation - probe orientation) and template center distance (current orientation - template center) and their interaction (Figure 1C). This model was preferred to a null model with only probe distance, χ²(2) = 10.53, p = 0.005. Examining the fixed effects revealed a main effect of template center distance, β = -0.01, CI = [-0.03 0.00], which reflected that when the template center was CW to the memory item, there were more frequent CW responses to the probe. We additionally found evidence for an interaction between template center distance and probe distances, β = 0.02, CI = [0.01, 0.03], such that the template center distance effect was attenuated for CW probes. We suspect that this directional attenuation may be due to response compatibility effects, since all responses were made with the right hand (e.g., Bauer & Miller, 1982).

Of note, we modeled the same effects in the unpredictable condition, and found no evidence for template distance effects, χ²(2) = 3.61, p = .164. These results imply that participants used the templates associated with the colors.

### The neural code of identical visual information depended on its predictability

Memory items were successfully decoded during encoding (Figure 2A), both for predictable ( 8–800 ms, M = 5.1×10^−3^, p<0.001) and unpredictable memory items (4 – 740 ms, M = 5.5×10^−3^, p<0.001). There were no significant differences between conditions. Similarly, both predictable (10–440 ms, M = 1.9×10^−3^, p < 0.001) and unpredictable (4–528 ms, M = 1.8×10^−3^, p < 0.001) memory items could successfully be decoded during memory maintenance following the impulse (Figure 2B), with no significant differences between conditions. Thus, despite the behavioral differences between conditions, the representations were decodable to a similar extent for both predicted and unpredicted memory items, which may suggest a functionally equivalent neural representation.

**Figure 2.**
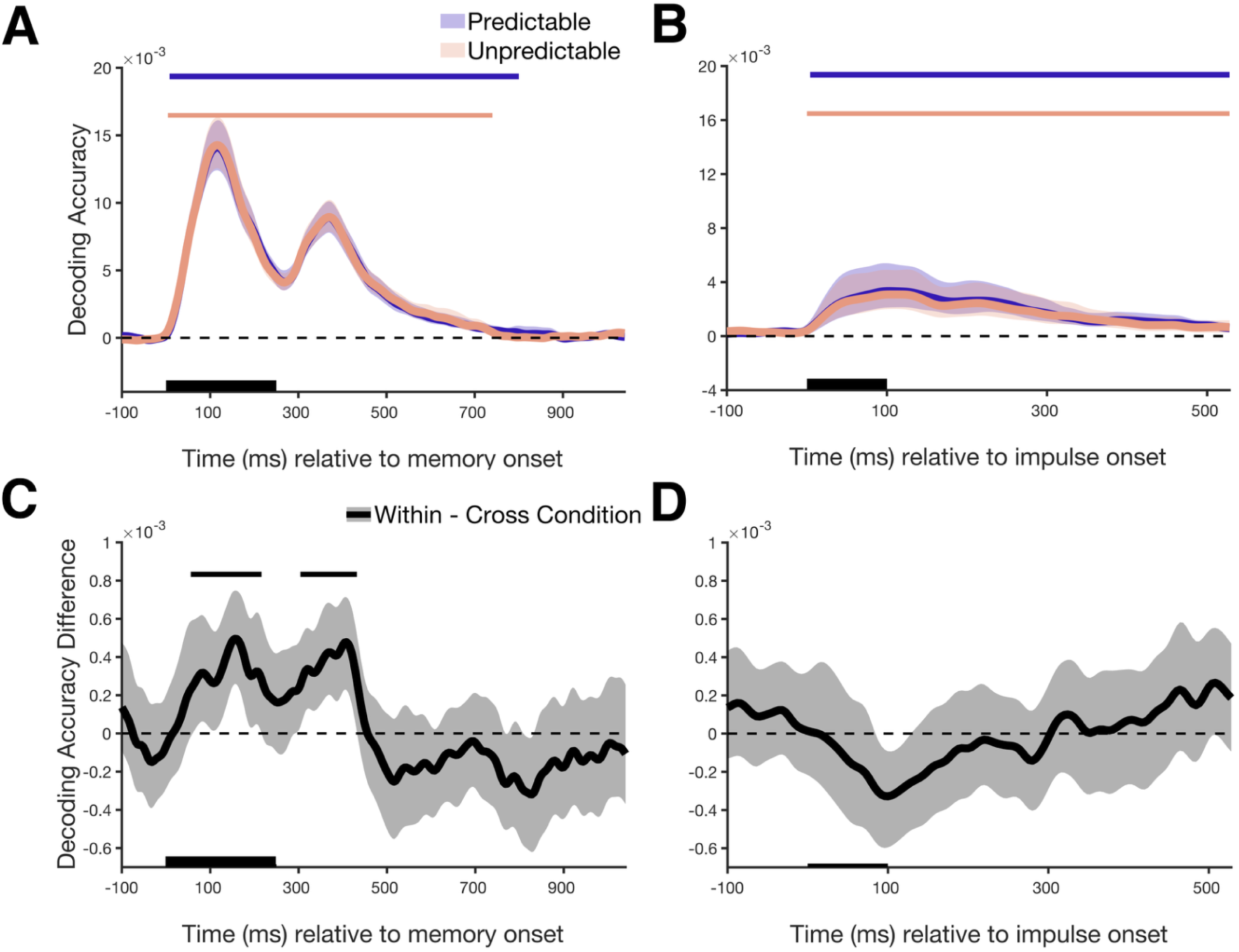
Time-course of decoding accuracy. **(A and B)** The solid lines show the mean decoding accuracy of the predictable and unpredictable condition. The shaded regions represent the 95% confidence interval. Horizontal lines represent the clusters with statistically significant decoding (*p <* 0.05, one-sided). The black bars on the x-axis indicate the time of memory item and impulse presentation, respectively. **(C and D)** Time-course of the cross – within condition means decoding accuracy. The horizontal line represents the clusters with statistically significant decoding (*p <* 0.05, one-sided).

However, a similar decoding accuracy does not necessarily mean the underlying neural code is the same. To assess this, we trained a classifier in one predictability condition and tested it on the other, then averaged both cross-condition decoding scores and compared this to the average within-condition decoding scores. This revealed that cross-condition decoding performed significantly worse than within-condition decoding (Figure 2C: 56–216 ms, average decoding accuracy = 1.11×10^−2^, *p* = 0.005; 304–432 ms, average decoding accuracy = 7.6×10^−3^, *p* = 0.008). This suggests that although memory items in both conditions were decodable to a similar extent, the underlying representation changed as a function of predictability, already during encoding.

Of note, cross-condition decoding in the maintenance epoch following the impulse stimulus did not differ from within-condition decoding (Figure 2D). Because of this, the following analyses focus on the representational differences during the encoding epoch.

To make sure that cue-specific representations did not factor into the observed differences, we explicitly sought to decode whether and to what extent we would be able to decode the cue category during the encoding of the memory item. Although this was possible, the clusters of significant color decoding did not coincide with significant cross-decoding differences (Supplementary Figure 1). Furthermore, we found evidence that cue category decoding scores did not correlate with cross-decoding differences across participants (see Supplementary Text 1). Taken together, we conclude that the predictions instill a change in the neural representation of orientation, independent of concurrently maintained cues.

### Representational similarity among items during encoding is increased when items are predictable

We next investigated how the neural representations of orientations were affected by predictability. To this end, we divided orientations into 12 orientation bins, and computed the average Mahalanobis distances between them to form Representational Dissimilarity Matrices (RDMs; Figure 3A). We realigned the bins across participants such that bins at -45 and +45° reflected the two template centers, intended to uncover potential structural patterns relating to the prediction templates.

**Figure 3.**
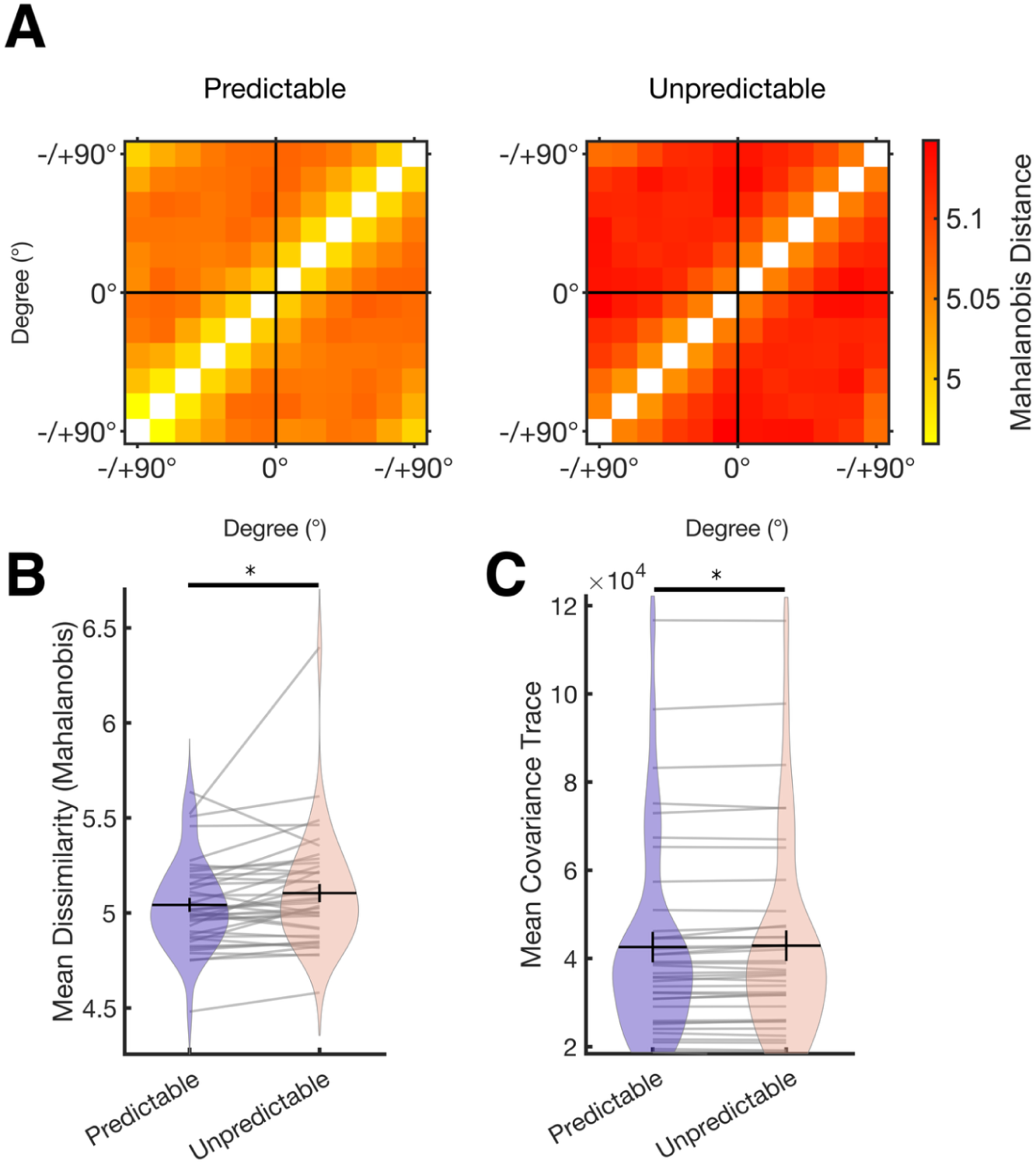
Representational similarity analyses and covariance comparisons. **(A)** The representational dissimilarity matrices (RDMs) per predictability condition. Each square of the RDMs represents an orientation bin. Colors of the squares represent the measured Mahalanobis distance between compared bins. **(B)** Violin plot of the RDM means per condition. Gray lines depict individual participant means. **(C)** Violin plot of the mean covariance matrix trace per condition. Gray lines depict individual participant means. (* p < 0.05)

A pairwise t-test comparing the RDMs (predictable vs. unpredictable) showed that Mahalanobis distances among orientation bins in the predictable condition were smaller than those in the unpredictable condition, t(39) = -2.32, p = 0.02 (Figure 3B), indicating that representations were more similar (i.e., less discriminable) when memory items were predictable. Next, we aimed to understand the ways in which predictability increased the representations’ similarity. Since Mahalanobis distance is defined by structural distance and covariance, we evaluated both these measures separately.

### Predictability reduced neural variance

To assess the variance in the data, we calculated the traces of the covariance matrices used in computing the RDMs. Pairwise comparison showed that the predictable condition had less variance than the unpredictable condition, t(39) = -2.33, p = 0.02 (Figure 3C)^1^. These results suggest that predictability reduced the neural variance of the information at encoding. Although decreased variance in the predictable condition is largely in line with previous results on prediction, it seemingly runs counter to the decreased Mahalanobis distances found in this condition (Figure 3A). To better understand how predictability changes neural representations, we next turn to Euclidian distance metrics unweighted by variance.

### The neural representational space of stimulus orientation shrinks when it is predictable

To assess how prediction structurally affected neural representation of orientation, other than affecting variance, we recalculated the RDMs with Euclidean distances, which unlike Mahalanobis distance do not factor in variance (Figure 4A). Although it was not significant, we again observed that the predictable condition RDM revealed higher similarity between orientation bins compared to the unpredictable condition, t(39) *=* -1.74, p = 0.09 (Figure 4B). Importantly, the absence of a significant difference between average RDMs does not mean the representational pattern it contains is identical. Therefore, we subsequently analyzed whether the patterns within RDMs significantly differed. For this, we plotted the data in a multidimensional space (as in Wolff et al., 2020; Figure 5A, left panel). Confirming our finding that the distances were smaller for predictable information, the plot showed a trend for the layout of the information being shrunk for predictable compared to unpredictable conditions. Based on this, we further hypothesized that the layout of the representations would have a significant spatial separation across conditions (Figure 5A, middle panel). To test this, we compared the bins that belong to the same condition (within condition distance) and different conditions (cross-condition distances, see Methods). Pairwise comparison showed that the average within condition bin distances were significantly smaller than cross-condition distances, t(39) = -10.12, p < 0.001, reflecting a separation of the representation across the predictability conditions (Figure 5A, right panel). These findings show that the geometry of the neural representations was indeed affected by predictability.

**Figure 4.**
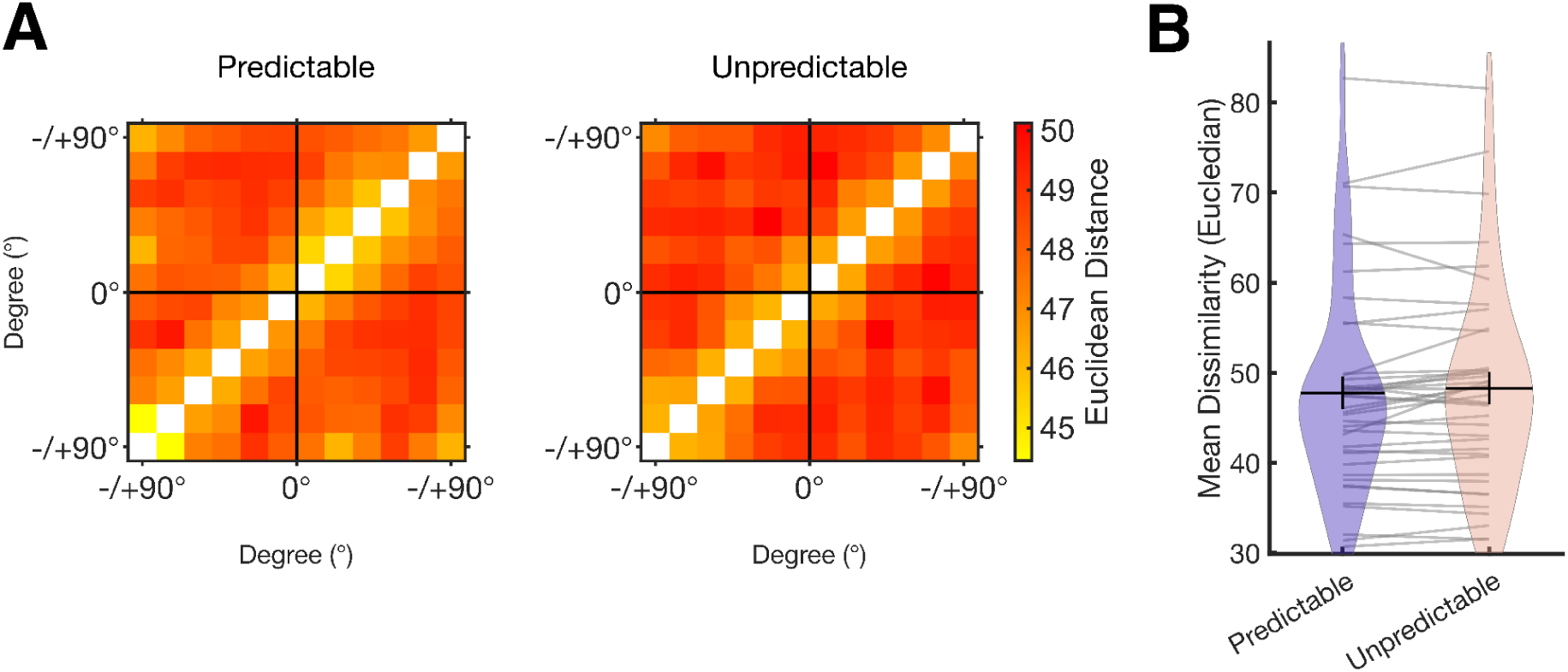
Representational similarity analyses with Euclidean distances. **(A)** The representational dissimilarity matrices (RDMs) per predictability condition. Each square of the RDMs represents an orientation bin. Colors of the squares represent the measured Euclidean distance between compared bins. **(B)** Violin plot of the RDM means per condition. Gray lines depict individual participant means.

**Figure 5.**
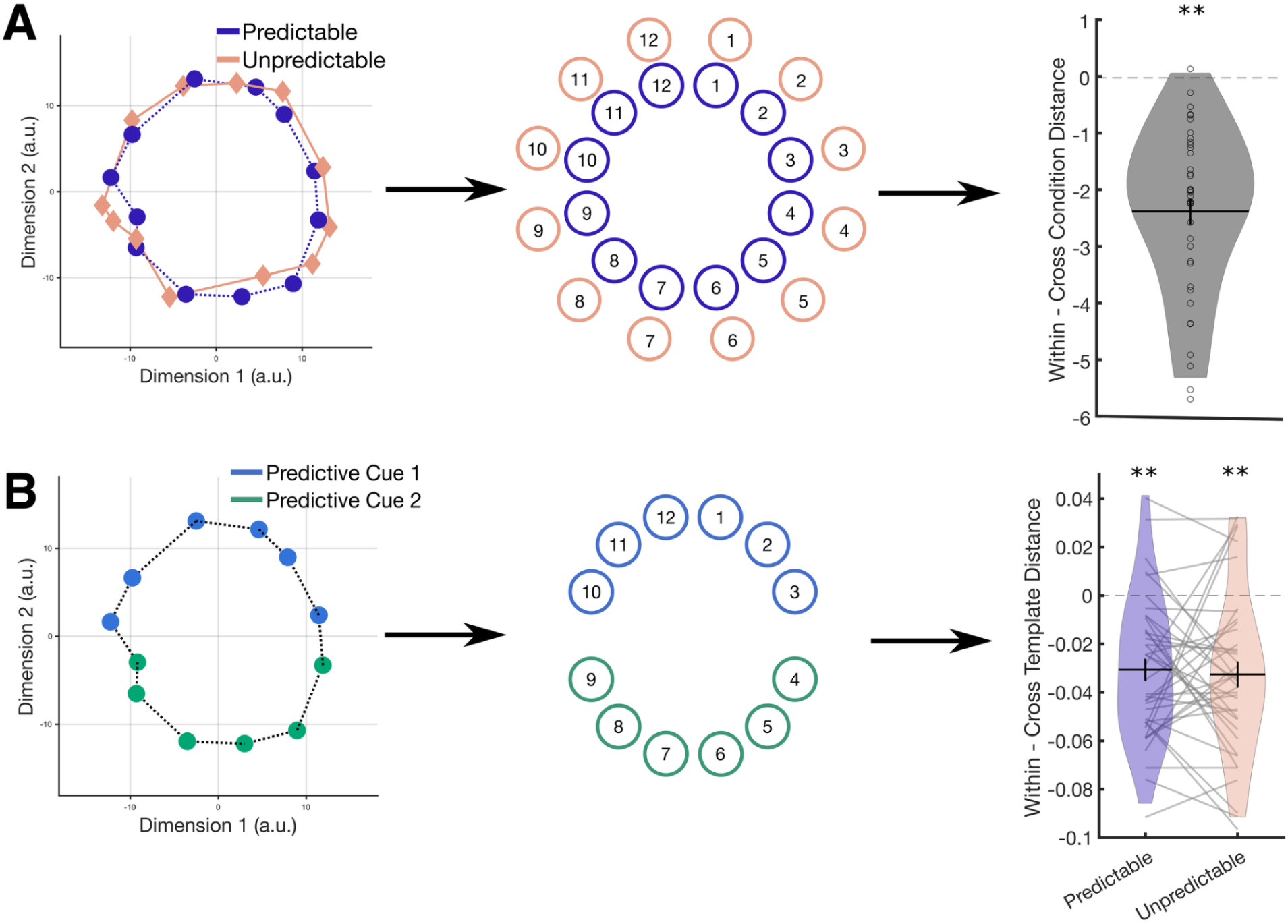
Assessment of the structural geometry of the neural representations. Figures depicting the observed (left panel), hypothesized geometry of the neural representations on a two-dimensional space (middle panel) and statistical testing of the hypothesized geometry defined by the within – cross condition (or template) bin distances (right panel, ** p < 0.001), for **(A)** cross-condition and **(B)** cross-template comparisons. Colored circles or diamonds represent the orientation bins.

### Neural representations are pulled towards the prediction templates

We investigated the effects of predictive templates on the shape of the representational structure by plotting the data in a multidimensional space (Figure 5B, left panel). The plot revealed a slightly oval-shaped layout of the information based on the template rather than a circle. Based on this, we hypothesized that the distances between bins that belonged to the same template (within-template distance) would be smaller compared to those belonging to different templates (cross template distances, see Methods, Figure 5B, middle panel). A pairwise comparison revealed significantly smaller distances within the template than across, t(39) = -6.61, p < 0.001 (Figure 5B, right panel). The result suggests that the representational layout indeed formed an oval rather than a circle, stretching the representation in line with the templates (Figure 5C).

### Irrelevant predictions stretch the unpredictable information

Because there was no predictive template in the unpredictable condition, we expected that the unpredictable condition would have no representational bias, in line with behavior. Regardless, we re-ran the analysis above for trials in the unpredictable condition, computing within and cross template distances. Interestingly, we found that within– and across–template distances were significantly different in the unpredictable condition as well, t(39) = -6.07, p < 0.001 (Figure 5D), and this effect was not significantly different across predictable and unpredictable conditions t(39) = 0.35, p = 0.726. This suggests that representations in the unpredictable condition were affected by the templates associated with the predictable condition. This could mean that, although suboptimal, the participants applied or compared the predictions they learned in the predictable condition to the unpredictable condition as well. We elaborate more on this in the Discussion section.

## Discussion

To investigate how diffuse predictability impacts neural representations in WM, we recorded EEG while participants judged whether a probe grating was rotated clockwise or counterclockwise relative to a predictable or unpredictable memory item. Memory item predictability was manipulated using cues, which were either nonpredictive or indicated a 90° range for the upcoming memory item orientation. We observed that predictive cues improved accuracy, and biased responses to align with the prediction. EEG data revealed that the neural representation for memory items differed between conditions during encoding, but not during maintenance, suggesting that representations at memory encoding changed depending on predictability. We showed that this representational change comprises reduced variance in the neural responses, and a shrunk representational spread for predictable information, alongside a bias incurred by the prediction templates. These findings illustrate how diffuse predictions reshape information encoding in WM.

Consistent with earlier work (Agres et al., 2018; Beck et al., 2014; Bollinger et al., 2010; Brady et al., 2009; Giesbrecht et al., 2006; Katzner et al., 2012; Kok et al., 2012; Padgaonkar et al., 2017; Ravizza, 2025), we observed a performance benefit for predictable information. Behavioral benefits were small, which is presumably due to the imprecise diffuse predictions offered by the cues, and due to the low difficulty of the single-item memory task. Nevertheless, participants’ responses were systematically biased by the predictions, revealing that they utilized the predictions, and that these predictions reshaped representations at memory encoding.

Predictable and unpredictable orientations were decodable from EEG to a similar extent. Importantly, cross-condition decoding nevertheless revealed a difference in their neural code. According to the Bayesian integration model, our brain combines prior knowledge and new sensory input (Friston, 2005, 2010; Körding & Wolpert, 2004; Lee & Mumford, 2003; Rao & Ballard, 1999). Our results directly reflect such integration, by showing that predictions altered the neural representation of incoming visual information. For precise predictions, such modulation has previously been found in the form of sharpening the neural representation (Yan et al., 2023; González-García & He, 2021; Kok et al., 2012). Our results on diffuse predictions, however, show that predictions shrunk the representational layout compared to unpredictable orientations, suggesting higher similarity between representations. This reveals a discrepancy between diffuse and precise predictions. Sharpening the neural representation of predicted information through precise predictions may benefit encoding processes when a single outcome is expected. In contexts where diffuse predictions are utilized, representations do not require strict dissociation within the cued information. In fact, increased similarity within the predicted range may be an efficient strategy when there are multiple possible outcomes with equal likelihood, as fine-grained dissociations would not provide additional advantage. We found that predicted information was represented with decreased neural variance, potentially through diffuse predictions stabilizing the neural code. This can be due to priors facilitating preparation of the visual processing for predicted information (Samaha et al., 2015). Extensive literature on prediction suggests that top-down control mechanisms influence perception based on internal priors (Friston, 2005, 2010; Körding & Wolpert, 2004; Lee & Mumford, 2003). An outcome of such preparation could be stabilizing the neural responses to the visual processing of the predicted information. Taking both reduced variance and shrunk representational distances for predictable information into account, our results suggest that diffuse predictions supported a robust and stable information structure that did not prioritize representational separability within the predicted template.

In line with the behavioral bias towards the prediction templates, neural representations were also pulled in towards the center of prediction templates. Interestingly, there was no behavioral effect of templates in the unpredictable condition, yet neurally we observed a similar representational bias. A possible explanation for this could be that learned predictions are retrieved in unpredictable situations even when they are not suitable. The brain is often described as a ‘prediction machine’ that continuously attempts to match the sensory input with generated predictive models (Clark, 2013). Because the brain inherently resists unpredictability (Peters et al., 2017), it might automatically attempt to evaluate learned predictions to novel unpredictable contexts, even when these are not applicable. Conversely, the observed neural bias could reflect an involuntary retrieval of the predictive template in response to presented information. Previous research showed that a stimulus can trigger incidental retrieval of its associated information (Kompus et al., 2011; Kuhl et al., 2013; Stark & Okado, 2003), and therefore presenting the unpredictable information might have triggered reactivation of the closest matching prediction template. Alternatively, the template that matched the unpredictable information could have been activated to be used as a categorical reference (Pereira Seabra et al., 2025; Yan et al., 2023). Whichever was the cause for neural reactivation of the template, it did not affect later responses to the probe, which sets it apart from the role of the templates in the predictable condition.

Unlike at encoding, we did not observe a neural code change during memory maintenance. A contributing factor to this may be that the neural signal in response to the impulse was rather weak compared to the response at encoding. This would make it difficult to identify clear differences between conditions. Seeing as we found a neural modulation during encoding, and a behavioral effect during the probe, this might suggest that the role of diffuse predictions is reduced or absent during maintenance. Nevertheless, the lack of effect at maintenance does not warrant space for interpretation.

Overall, our findings indicate that diffuse predictions stabilize and reshape the neural code during stimulus encoding, by decreasing variance and biasing the representational structure in line with the predictions. The observed neural and behavioral biases reflect an integration of prior and sensory evidence, suggesting that the brain represents a ‘best guess’ of sensory input rather than faithfully reflecting our environment.

## Methods

### Participants

Forty-seven volunteers (36 female, mean age: 23.4, range: 18-31) participated in the experiment in return for either course credit or monetary compensation. Four participants were excluded due to low performance (average performance < 60%), and three were excluded due to poor EEG data quality (see EEG acquisition and preprocessing). All participants were informed about the experimental task, EEG data collection procedure, and data sharing procedures, and provided written informed consent prior to experiment. This study was conducted in accordance with the Declaration of Helsinki, under the ethics approval of the Leibniz Research Centre for Working Environment and Human Factors Committee.

### Stimuli and procedure

The experiment was conducted while participants sat in a dimly lit soundproof experimental chamber in front of a 19-inch CRT monitor screen with 100 Hz refresh rate and 1,280 by 1,024 pixels resolution. The viewing distance was 80 cm. The task was coded and performed using MATLAB (R2024a) with Psychtoolbox (Version 3.0.19). Memory items and probes consisted of sine wave gratings with a spatial frequency of 0.65 cycles per degree and a circular edge. Gratings were presented centrally with 20% contrast and a diameter of 6.69° of visual angle. The phase of the gratings was randomized across trials. The cues were colored dots of 0.73°, presented at the center of the screen in red (RGB = [255, 0, 0]), green (RGB = [0, 255, 0]) and blue (RGB = [255, 0, 0]) colors. A bullseye stimulus with the same size and spatial frequency of the memory items but with sharp black-white transitions was used as an impulse stimulus. The fixation dot was black (RGB: [0, 0, 0]), 0.73°, and was presented at the center of the screen throughout the trial when there were no other stimuli displayed. The background of the experimental task was set to gray (RGB = [128, 128, 128]).

Across participants, colors were assigned to be predictive or unpredictive, in a counterbalanced fashion. The two ranges of orientation cued by the two predictive colors were randomly determined per participant. Cues were 100% valid. Before the experiment started, participants were explicitly shown how the different colors related to possible upcoming memory items. The instruction was repeated until it was confirmed by the experimenter that participants fully understood the associations. Next, the participant performed the experimental task. The trial procedure is depicted in Figure 1A. Each trial began with a randomly sampled intertrial interval (1200-1500 ms), during which only a fixation dot was on screen. This fixation dot was presented throughout each delay interval. Next, the cue was presented for 250 ms, followed by a delay interval of 900 ms. Then, the memory item was shown for 250 ms, followed by another delay interval of 900 ms. An impulse display, which was on screen for 100 ms was followed by another delay interval of 500 ms. The memory probe was an orientation grating that deviated from the memory item (±5, 10, 16, 24, 26°), and was displayed for 250 ms. This was followed by a maximum of 1000 ms response interval. Participants judged whether the probe item on the screen was CW or CCW compared to the memorized item, by pressing the right or left arrow key respectively. Following each response, a happy or sad smiley face was presented for 150 ms, as a feedback. To make sure participants were attending to the cue, at the end of 20% of the trials a cue probe appeared, testing participants on which color had been presented on this trial. The three cue colors were displayed horizontally for 2000 ms in a random order, and participants responded with the left, down or right arrow key.

The experiment consisted of 1800 trials that were divided into 20 blocks. Participants took self-paced breaks between each block. During breaks, they saw their performance on the previous block and a motivating sentence. During a block, trials were presented one after another until the block ended.

### Analysis of behavioral data

Behavioral data was analyzed with a binomial GLMM with a Cauchy link function. To assess whether predictability of the memory item had an effect on behavior, we fit a GLMM to predict the probability of a correct response, with condition (predictable vs. unpredictable) as a fixed effect and by-subject random intercepts. To assess response biases relative to the template center, we fit another GLMM predicting the probability of giving a clockwise response. Fixed effects were template-center distance, probe distance and their interaction. This model included by-subject random intercepts and random slopes for probe distances, and was fit to data from predictable and unpredictable conditions separately. Model comparisons were performed using likelihood ratio tests.

### EEG acquisition and preprocessing

EEG was recorded using 64 Ag/AgCI passive electrodes (Easycap Gmbh, Herrsching, Germany) laid out in accordance with the 10-5 system and with 1000 Hz sampling rate. Data was recorded using NeurOne Tesla AC amplifiers (Bittium Biosignals Ltd, Kuopio, Finland) with a 250 Hz low-pass filter. The FCz electrode was set as the reference and AFz was set as the ground. Horizontal and vertical eye movements were tracked by electrodes that were placed above and below the right eye and the outer edges of both eyes, subtracting horizontally and vertically positioned electrodes to create electrooculography channels. The impedances of all electrodes were kept below 10 kΩ at the beginning of the experiment, and checked again after ten blocks.

Preprocessing of the data was done using the EEGLAB (2024.2; Delorme & Makeig, 2004) toolbox in MATLAB (R2024a). EEG data was down-sampled to 500 Hz and band-pass filtered between 0.1 and 40 Hz. Visually identified noisy channels were replaced using spherical interpolation and the data was re-referenced to the global average of all electrodes. Then, data was epoched -300 to 1100 ms from the memory item onset and -300 to 600 ms from the impulse onset. Independent component analysis (ICA) was used to identify and remove eye blink artefacts from the data. After the data was converted to the FieldTrip format (Oostenveld et al., 2011), ‘ft_rejectvisual.m’ was used to visually inspect and reject epochs with high noise. For this, we used the ‘summary’ method to exclude epochs with either high signal variance (>1500 µV), exceeding absolute z-values of 6 with the mean and standard deviation computed over all time points and epochs, and excessive absolute amplitudes (>120 µV) within each channel. Participants with more than 30% of the memory item or impulse epochs rejected were removed from the subsequent EEG and behavioral analysis due to low overall signal quality. The EEG analyses described below were performed over a set of 17 parietal-occipital electrodes, as used in previous WM decoding studies (Wolff et al., 2017; P7, P5, P3, P1, Pz, P2, P4, P6, P8, PO7, PO3, POz, PO4, PO8, O1, Oz, O2).

### Time-course decoding of the orientations

We decoded the memorized orientations following Kandemir et al. (2024). We decoded from the memory item and impulse epochs separately for predictable and unpredictable conditions. For each time point, the data was dynamically baselined using a sliding window approach. This was done by computing the average voltage in a 100 ms time window for each channel and epoch, centered on each time point. Then, this average was subtracted from all samples within the time window., the EEG data for each time point was restructured into channel ⨉ time feature vectors per trial.

For decoding, each epoch was given one of twelve classification labels, assigned by dividing presented orientation angles into twelve equal bins. Subsequently, epochs were divided across eight folds for cross validation, equalizing the trial count per orientation bin in all folds. Using each fold as a test set once, and the remaining folds as training sets, a covariance matrix was computed across all trials in the training set, as well as multivariate averages across trials per bin. In the test set, the Mahalanobis distance was computed between each trial and these bin centers at each time point. The resulting distances were averaged per bin in the test set, centered per bin, sign-reversed, and mean-centered. Finally, distances were cosine-weighted, to produce a decoding score reflecting the amplitude of a cosine-shaped tuning curve at each time point.

This procedure was repeated across 100 repetitions to minimize sensitivity to the specific definition of the folds. The orientation bins were redefined eight times by phase-shifting their offset in eight steps, to minimize sensitivity to exact bin placements. The decoding scores across these 100 ⨉ 8 repetitions were averaged to yield a single decoding time course per condition per participant.

Cross-condition decoding followed the same approach, but training data from one condition was used to assess decoding for testing data from the other condition. The two resulting cross-condition decoding results were averaged.

### Representational similarity analysis

Representational similarity analysis was conducted in order to assess the representational pattern among orientations (as in Kandemir et al., 2024)^2^. The EEG data was prepared as in the decoding analysis. Representational dissimilarity matrices (RDMs) were calculated over 12 orientation bins spanning from -90° to +90° per condition. The orientation bins were re-centered such that -45° and +45° corresponded to the template centers. Rather than decoding per time point, a single distance measure was computed using covariance matrix estimates across all time points in the epoch. To control for differences in trial counts across bins, trials from all bins were randomly subsampled to match the bin with the minimum number of trials. Then the trials for each bin were averaged. Mahalanobis distances between all twelve bin averages were calculated across 100 repetitions and averaged to minimize sampling bias. The resulting distances defined the RDMs.

We additionally investigated structural change in the representation across conditions, without reweighting via covariance. For this, we computed RDMs for both conditions reflecting Euclidean distances between orientation bins.

### Representational dissimilarity across conditions and across templates

For within - cross condition comparison we computed the representational distance of a bin to its adjacent bin within the same condition or in the other condition. Thus, the adjacent within and across bins corresponded to the same angular difference in orientation, with the only difference being the conditions the bins belong to. We took the difference between within and cross-condition bin distances per participant.

For within - cross template comparison we computed the representational distance of all bin combinations that belong to the same template or different templates. These bin combinations corresponded to the same angular difference in orientation with the only difference being the templates the bins belong to. We took the difference between within and cross template bin distances and averaged them per participant.

### Statistical testing

We statistically assessed EEG decoding scores using cluster-based permutation testing. A null distribution was generated by randomly flipping the sign of individual time courses 10,000 times and recomputing the mean at each time point. For each permutation, a series of neighboring time points where the mean exceeded the permutation-quantile threshold (0.05) were grouped into adjacent clusters. Each cluster was scored with the sum of its samplewise amplitudes. Then, observed cluster sums were compared against the null distribution of cluster sums to determine significance (p < 0.05).

## Data availability

The raw EEG data^3^ is available on the OpenNeuro repository, https://doi.org/10.18112/openneuro.ds007431.v1.0.0. The analysis results and the behavioral data is available on the Open Science Framework (OSF) repository, https://osf.io/8evwh/.

## Code availability

The custom scripts used for preprocessing the EEG data and subsequent analyses are available on the OSF repository, https://osf.io/8evwh/.

## Declaration of competing interests

Authors declare no competing interests.

## Acknowledgement

We thank Rama Alnabusy for her efforts in data collection.

## Author contributions

Conceptualization, N.A., Ș.Ö., W.K., D.S, and E.G.A.; methodology, N.A., Ș.Ö., W.K., D.S, and E.G.A.; formal analysis, N.A.; software, N.A. (developed), Ș.Ö. (contributed), and W.K. (supervised); visualization, N.A.; writing (original draft), N.A.; writing (review and editing), W.K., D.S, and E.G.A.; funding acquisition, D.S. and E.G.A.; supervision, W.K., D.S. and E.G.A.

## Footnotes

1 Mahalanobis distances in Figure 3C are computed using a Ledoit-Wolf shrinkage estimator to estimate the covariance matrix. Comparing distances using raw covariance matrices without shrinkage gave identical comparison statistics, t(39) = -2.33, p = 0.02.

2 For the corresponding script in Kandemir et al., (2024) please see https://github.com/mijowolff/veridical-and-transformed-representations-in-wm/blob/main/Fig5_impulse2_RSA_spatiotemporal.m

3 The participant numbers in the raw dataset have been randomly reassigned to prevent participant identifiability.

## Notes

### Competing Interest Statement

The authors have declared no competing interest.

