## Supplementary Text 1 for "Diffuse predictions stabilize and reshape the neural code during working memory encoding"

### Supplementary Materials

**Supplementary Text 1.** A possible explanation for the reduced cross-condition decoding during memory item display could stem from the lingering effect of the cue at that time interval. Even though it is not possible to differentiate the effect of the cue color from the predictive template that was associated with the cue, we aimed to rule out both possibilities to prove that what is changed is indeed the representation of the memory item itself. For this, first we assessed whether the cue was decodable during memory item display. We found that, although noisy, the cue categorization was significant over a few clusters (Supplementary Figure 1A). Next, we assessed whether the cue categorization was significant over the clusters where cross-condition decoding was significantly reduced. We found that for the early cluster cue categorization was not possible ($BF_{10}$ = 0.195), however for the later cluster it was ($BF_{10}$ = 4.180, Supplementary Figure 1B). Finally, we addressed whether the reduction in cross-condition generalization correlated with cue categorization, and found evidence against such a correlation for both clusters (cluster 1, r = -0.186, $BF_{10}$ = 0.371; cluster 2, r = 0.137, $BF_{10}$ = 0.277, Supplementary Figure 1C). This suggests that even though there is a lingering representation of the cue, it did not show a relation to the time-course of the representational change during encoding.
