## Supplementary Figure 1 for "Diffuse predictions stabilize and reshape the neural code during working memory encoding"

### Supplementary Materials

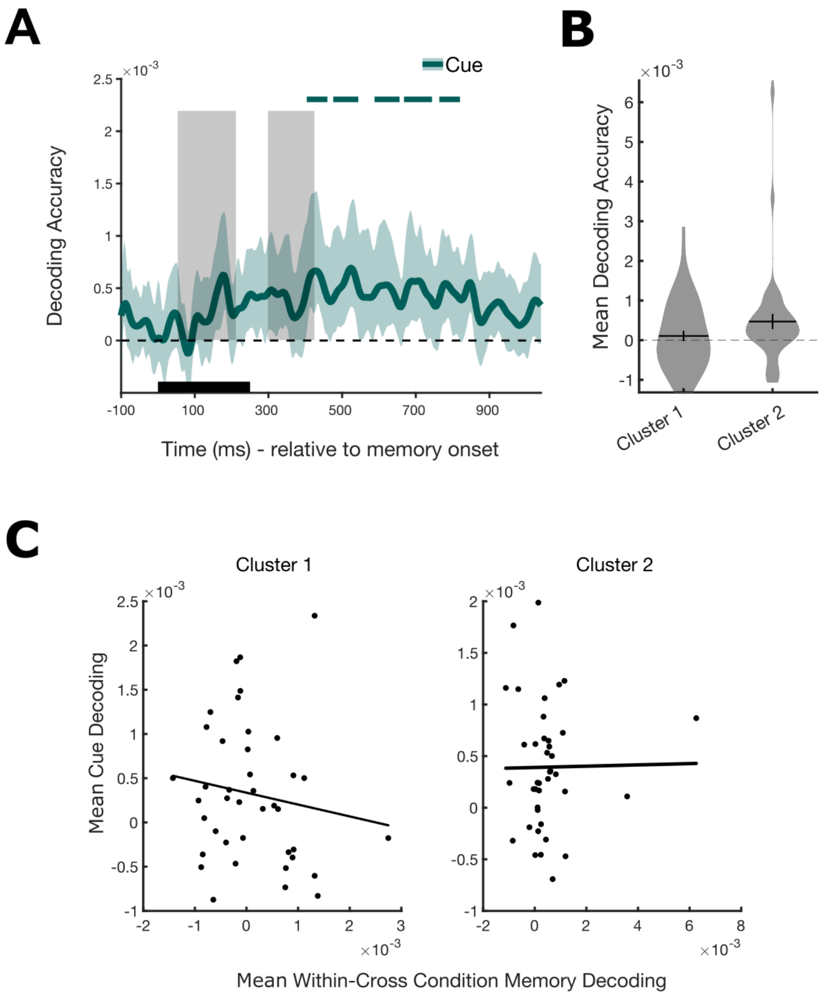


**Supplementary Figure 1. Decoding the cue at the memory item display. (A)** The solid line shows the time-course decoding accuracy for the cue averaged over cues and the shaded ribbons surrounding the line represent the 95% confidence interval. The horizontal lines on the top of the figure indicate the clusters with statistically significant cue decoding ($p < 0.05$, one-sided). The shaded gray boxes depict the two clusters where within – cross condition decoding of the memory item was significantly different from zero (see Figure 2C). **(B)** Violin plot of the mean decoding accuracy of the cue for each cluster. **(C)** Scatter plots showing the average within – cross condition decoding of the memory item and cue decoding per participant. The left panel represents the first cluster where within – cross condition decoding was significantly different from zero, and the right panel represents the second cluster.
